# Targeting platelet GPVI or GPIIb/IIIa only minimally affect acute inflammation and cardiac repair in experimental permanent myocardial infarction

**DOI:** 10.1101/2025.10.22.683606

**Authors:** Anna Rizakou, Lukas Johannes Weiss, Ecem T. Sakalli, Giuseppe Rizzo, Sarah Beck, Philipp Burkard, Stefano Navarro, Annabelle Rosa, Sourish Reddy Bandi, Marie Piollet, Vanessa Göb, Tobias Krammer, Antoine-Emmanuel Saliba, David Stegner, Alma Zernecke, Clément Cochain, Bernhard Nieswandt

## Abstract

**Background:** Beyond their role in hemostasis, platelets are recognized as key regulators of inflammatory responses in ischemic diseases, including cardiac ischemia/reperfusion (I/R) injury with key roles of platelet membrane glycoproteins (GP)VI and IIb/IIIa. However, whether platelet-driven thrombo-inflammatory pathways affect acute inflammation and cardiac repair processes in permanent, non-reperfused myocardial infarction (MI) is unknown.

**Methods:** We targeted GPVI and GPIIb/IIIa in experimental permanent MI in mice. Cardiac, bone marrow, and blood innate immune responses were evaluated by flow cytometry and single-cell RNA-sequencing. Survival and cardiac repair were assessed over the inflammatory and scar formation phase, until day 10 after permanent MI.

**Results:** Platelet GPVI immunodepletion by injection of the anti-GPVI antibody JAQ1 did not affect levels of neutrophil or monocyte subsets (Ly6C^hi^ and Ly6C^low^) in the bone marrow and blood, and did not alter accumulation of monocytes, macrophage subsets (defined by expression of MHCII and TIM4), or neutrophil subsets (SiglecF^hi/low^) in the infarcted heart on day 4. GPVI depletion only had a minimal effect on cardiac repair, slightly decreasing interstitial fibrosis in the infarct border zone on day 10. Four days after MI, GPIIb/IIIa inhibition by JON/A-F(ab′)_2_ had no effect on cardiac or systemic innate immune cell levels as measured by flow cytometry and did not affect composition and transcriptomic profile of the cardiac immune infiltrate as revealed by single-cell RNA-sequencing. GPIIb/IIIa inhibition did not improve cardiac remodeling, and was even associated with an increased mortality rate over 10 days post-MI.

**Conclusion:** Targeting the GPVI-GPIIb/IIIa axis only had minor effects on post-MI inflammatory responses and cardiac wound healing in permanent myocardial ischemia. Our findings demonstrate that the therapeutic benefits of inhibiting platelet-driven thrombo-inflammation are particularly relevant in the subacute reperfusion phase.

## Introduction

Dysregulated platelet activation and thrombus formation are the most frequent causes of myocardial infarction (MI). Early recanalization of the infarcted vessel and prevention of recurrent thrombosis are key objectives in cardiovascular (pharmaco-)therapy. After reperfusion, immune cells infiltrate the heart, mediating wound healing and scar formation during the reparative phase^1^, and long-term ventricular remodeling but also acute post-MI arrhythmias in the ischemia/reperfusion (I/R) process^2,3,4^. This post-MI inflammatory response is a systemic process involving activation of emergency myelopoiesis in the bone marrow, leading to the production of monocytes and neutrophils that are mobilized to the bloodstream and ultimately recruited to the heart where they contribute to local inflammation^5^. The mutual interplay of platelet and innate immune effector pathways, now commonly referred to as thrombo-inflammation^6,7^, is crucially involved in post-MI inflammatory responses^8,9^. Platelets express multiple surface receptors that are involved in the interaction with immune cells, including the immunoreceptor glycoprotein (GP) VI, and the integrin GPIIb/IIIa which upon conformational activation binds multiple ligands, including fibrin(ogen) and von Willebrand factor (vWF). In ischemia/reperfusion models of the heart and brain, depletion or blockade of GPVI and GPIIb/IIIa, respectively, were shown to reduce myocardial thrombo-inflammation and improve the outcome^10,11,12,13,14^. Novel antithrombotic agents targeting GPVI function have been developed, notably the humanized anti-GPVI antibody Fab-fragments EMA601 and glenzocimab^11,15^. While the pathogenic role of the GPVI-GPIIb/IIIa axis in thrombo-inflammation is well established in the context of reperfused ischemia of the heart^16^ or brain^10^, it is unclear whether this pathway affects local and systemic inflammatory responses and tissue repair processes in permanent myocardial ischemia. This model reflects the substantial proportion of MI patients without reperfusion and is usually used to assess wound healing and scar formation in the infarcted heart^17^.

Here, we aimed to investigate the effect of inhibiting thrombo-inflammation in permanent myocardial ischemia by GPVI immunodepletion and GPIIb/IIIa blockade. We comprehensively assessed local and systemic acute inflammation and cardiac repair after MI using flow cytometry and single-cell RNA-sequencing (scRNA-seq). In stark contrast to I/R, we show that targeting the GPVI-GPIIb/IIIa axis only had minor effects on post-MI inflammatory responses and cardiac wound healing in permanent myocardial ischemia. Our findings demonstrate that the therapeutic benefits of inhibiting thrombo-inflammation are of particular relevance in the setting of reperfusion.

## Results

### Targeting GPVI does not affect local and systemic acute inflammation after permanent MI

GPVI is the central activating platelet collagen/fibrin receptor. *Gp6* knockout (KO) mice exhibit protection against arterial thrombosis without an increased risk of bleeding, highlighting its potential as a target for antithrombotic therapy. Beyond its direct role in thrombosis and hemostasis, GPVI has also been identified as a key mediator of thrombo-inflammation^11,18^. Previous work has shown that antibody-mediated GPVI depletion reduces infarct size in cardiac I/R injury^10^. However, the mechanisms by which GPVI depletion improves the outcome are not fully understood. We hypothesized that GPVI depletion affects leukocyte-mediated remodeling post-MI. To test this, we assessed thrombo-inflammation following permanent coronary occlusion, as this model allows longitudinal studies of myocardial remodeling. We immunodepleted platelet GPVI by injection of the anti-GPVI monoclonal antibody (mAb) JAQ1^19^, with two injections performed 6 and 4 days prior to permanent ligation of the left anterior descending coronary artery (LAD) (**Figure 1a**). Systemic and cardiac inflammation were analyzed 4 days after MI by flow cytometry (see **Methods** for pre-gating strategies), at which time efficient GPVI depletion was confirmed (**Figure 1b**). In the bone marrow (**Figure 1c-d**) and blood (**Figure 1e-f**), GPVI depletion did not affect the levels of monocytes and their distribution among the Ly6C^hi^, Ly6C^int^ and Ly6C^low^ subsets. Cardiac infiltration of macrophages, monocytes and neutrophils was analyzed by flow cytometry (**Figure 1g-j**). Absolute cell counts and proportions of total CD11b^+^Ly6G^-^F4/80^+^Ly6C^low^ macrophages among CD45^+^ leukocytes were not altered by GPVI depletion, and the distribution of macrophage subsets defined by surface expression of MHCII and TIMD4^20,21^ (TIMD4^+^MHCII^+^ and TIMD4^+^MHCII^-^: tissue-resident macrophages; TIMD4^-^MHCII^+^: resident and recruited MHCII^+^ macrophages; TIMD4^-^MHCII^-^: recruited MHCII^-^ monocyte-derived macrophages) was unchanged (**Figure 1g-h**). Infiltration of CD11b^+^Ly6G^-^F4/80^low^Ly6C^high^ inflammatory monocytes was also unaffected by GPVI depletion (**Figure 1g,i**). Absolute cell counts and proportions of total CD11b^+^Ly6G^+^ neutrophils, and distribution of neutrophils among SiglecF^low^ and SiglecF^high^ subsets^22^ were not affected (**Figure 1j**). Altogether, these results indicate that loss of functional GPVI does not affect local and systemic inflammatory responses in the setting of permanent MI in mice.

**Figure 1:**
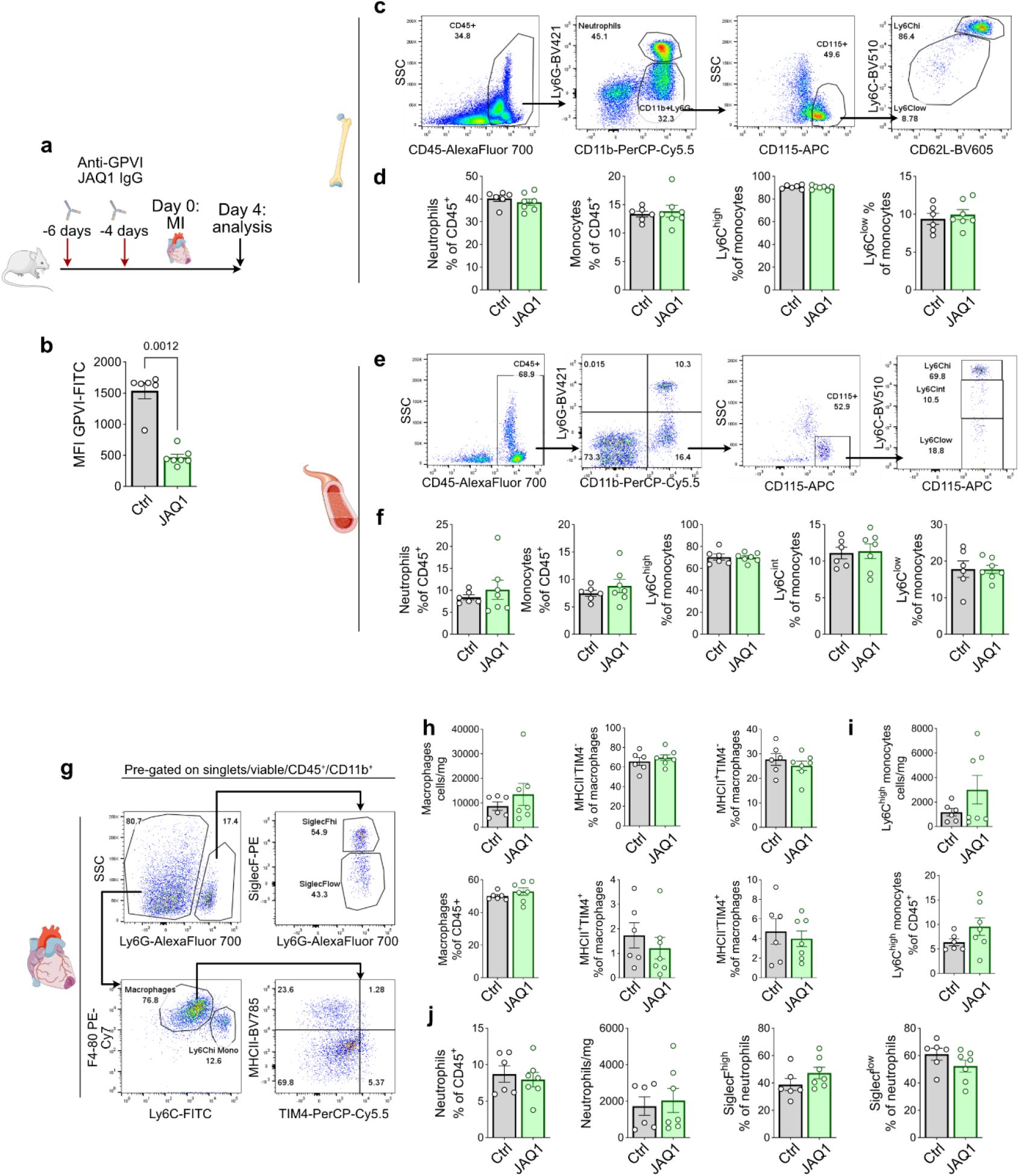
GPVI depletion does not affect acute inflammatory responses after permanent occlusion. **a)** experimental workflow; **b)** Platelet GPVI levels measured by flow cytometry; **c)** representative flow cytometry gating strategies (pre-gated on viable singlets) and **d)** quantitative analysis of bone marrow myeloid cells; **e)** representative flow cytometry gatings (pre-gated on viable singlets) and **f)** quantitative analysis of circulating myeloid cells; **g)** representative flow cytometry gatings of cardiac myeloid cells; **h-j)** quantitative analysis of **h)** cardiac macrophages and macrophage subsets, **i)** cardiac monocytes and **j)** cardiac neutrophils and neutrophil subsets in the heart; isotype control group, n=6; anti-GPVI (JAQ1) treated group, n=7. Data presented as mean ± SEM.

We next investigated the impact of platelet GPVI loss on cardiac repair after permanent MI. We focused our analysis on readouts related to scar formation over 10 days, a time point where acute inflammation has resolved and myofibroblast-mediated fibrotic scar formation has occurred^4^. We measured survival, infarct size, and collagen deposition in the infarct border zone. Mice were treated 6, 4 and 2 days before MI with JAQ1 to induce platelet GPVI depletion, which was maintained by a fourth injection on day 4 post-MI (**Figure 2a**). Platelet GPVI depletion did not significantly affect survival (**Figure 2b**) or infarct size (**Figure 2c-d**) but was associated with a small yet significant decrease in collagen deposition in the infarct border zone (Control: 14.44±1.99%; JAQ1: 11.0±1.53%; p=0.012; **Figure 2e-f**). These results suggest that loss of functional GPVI only slightly affects cardiac repair after permanent MI, with an effect limited to a small decrease in interstitial fibrosis in the border zone.

**Figure 2:**
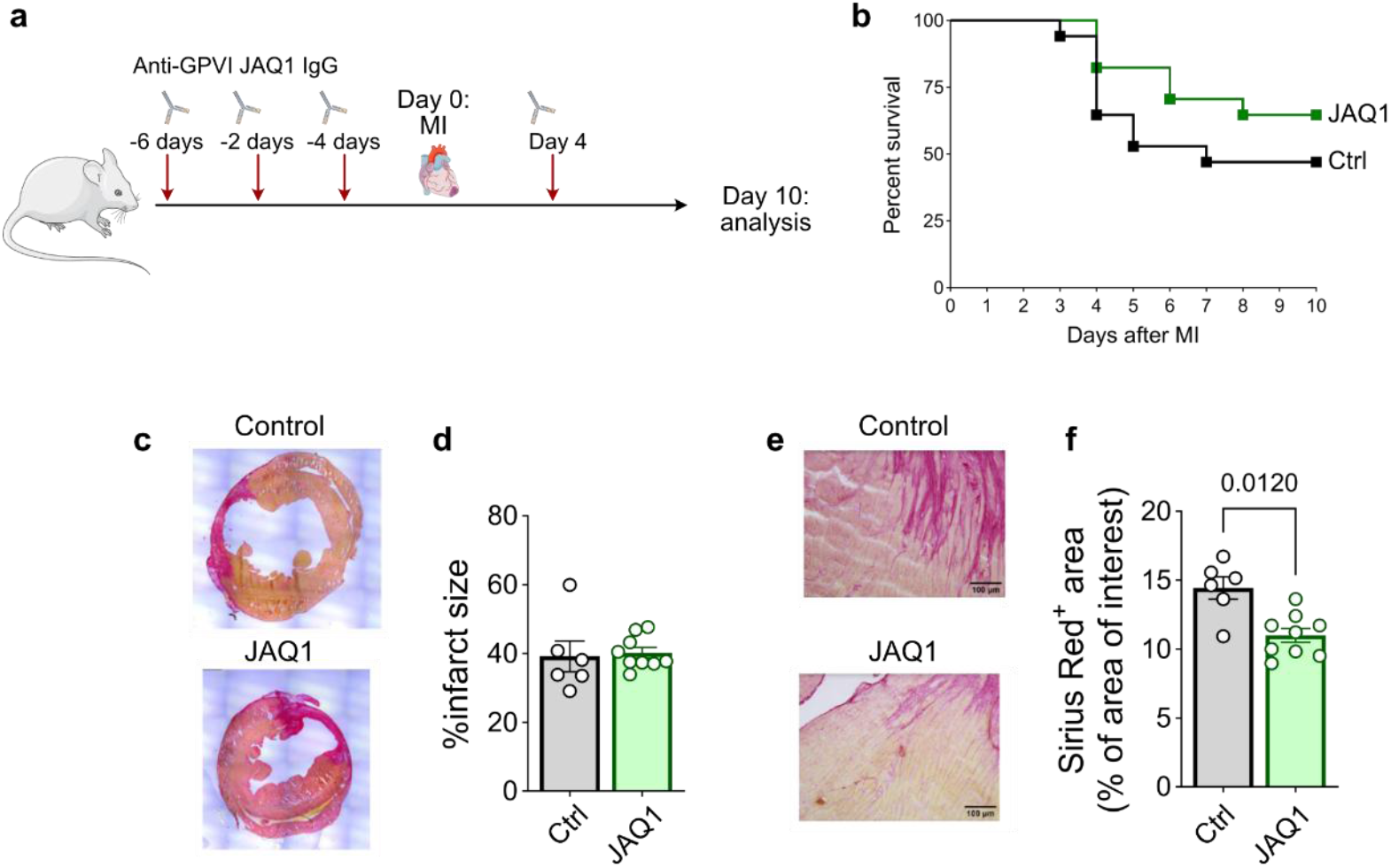
GPVI depletion minimally affects cardiac remodelling after permanent LAD occlusion. **a)** experimental workflow; **b)** Survival plot of isotype control (n=18) and anti-GPVI (JAQ1) treated (n=15) mice over 10 days after MI; **c)** Representative Picrosirius red staining of heart sections and **d)** quantitative analysis of infarct size at day 10 after MI. **e)** Representative staining of Picrosirius red heart sections and **f)** quantitative analysis of picrosirius red positive staining in the border zone. Scale bars: 100 μm. Results pooled from 3 experimental series. **c-f**n control mice n=6; JAQ1 treated mice n=9. Scale bars in panel e indicate 100 µm. Data presented as mean ± SEM. Panel b: log-rank (Mantel-Cox) test; panel d and f: Mann-Whitney test.

### Targeting GPIIb/IIIa minimally affects local and systemic acute inflammation after MI

Beyond immunoreceptors, platelets abundantly express adhesion receptors of the integrin class, which mediate the interaction with extracellular matrix (ECM) proteins and other macromolecular ligands. GPIIb/IIIa is the dominant platelet integrin which upon conformational activation in response to platelet agonists binding binds multiple ligands, including fibrin(ogen) and vWF. Functional GPIIb/IIIa is essential for platelet aggregation and thrombus formation, but the integrin has also been identified as a central mediator of thrombo-inflammation. Due to its critical role in these processes, GPIIb/IIIa is an established target of several clinically approved antithrombotic drugs such as eptifibatide or abciximab. We next investigated the effect of GPIIb/IIIa blockade on post-MI inflammation using the anti-GPIIb/IIIa antibody JON/A-F(ab′)_2_^23^. As GPIIb/IIIa blockade induces a strong hemostatic defect, mice were treated 1 hour and 2 days after permanent MI. The analysis was performed 4 days after LAD occlusion (**Figure 3a**). GPIIb/IIIa accessibility was blunted as detected by reduced JON/A antibody-binding by flow cytometry^23^ (**Figure 3b-c**). In the bone marrow, a slight but statistically significant increase in neutrophils, and decrease in monocyte proportions among total CD45^+^ leukocytes were observed. Within monocytes, a slight but significant shift from Ly6C^hi^ towards Ly6C^low^ monocytes was noted (**Figure 3d**). No differences in blood monocytes and neutrophils were observed (**Figure 3e**). Levels of cardiac macrophages, monocytes and neutrophils, and distribution of macrophages and neutrophils among subsets defined by flow cytometry were unaffected by JON/A-F(ab’)_2_ treatment (**Figure 3f-h**).

**Figure 3:**
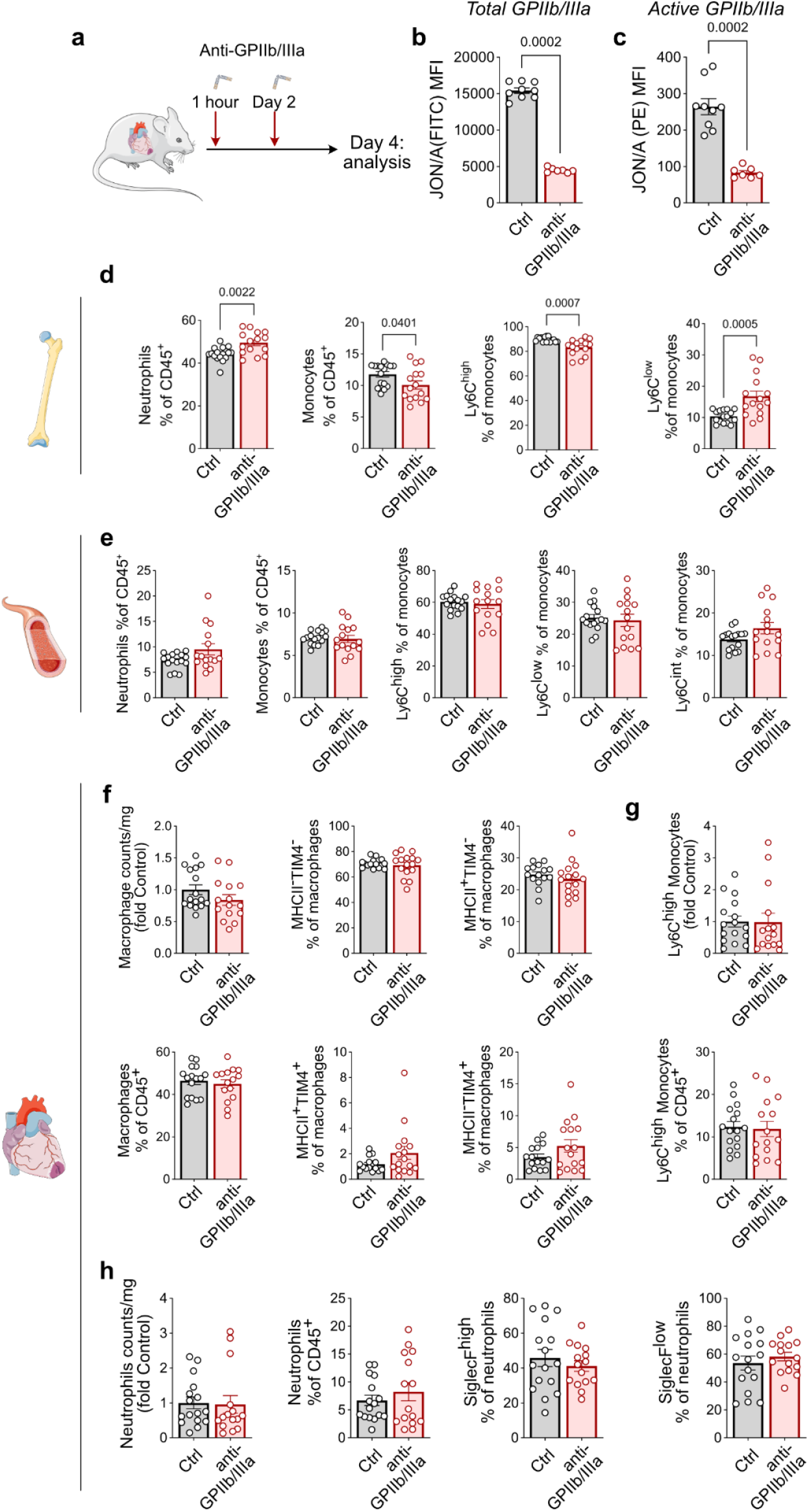
GPIIb/IIIa blockade and post-MI acute inflammatory responses. **a)** experimental workflow; **b)** platelet resting and **c)** activated GPIIb/IIIa levels measured by flow cytometry **d)** quantitative analysis of bone marrow myeloid cells; **e)** quantitative analysis of blood circulating myeloid cells **f-h)** quantitative analysis of **f)** cardiac macrophages and macrophage subsets, **g)** cardiac monocytes and **h)** cardiac neutrophils and neutrophil subsets in the heart; isotype control group, n=16; anti-GPIIb/IIIa (JON/A-F(ab’)_2_) treated group, n=15; results pooled from 2 independent experiments. Data presented as mean ± SEM.

### Targeting GPIIb/IIIa does not affect cardiac leukocyte composition: single-cell transcriptomic analyses

Flow cytometric analysis is inherently limited by the number of epitopes that can be simultaneously labeled, and by the lack of cell surface markers clearly defining some transcriptionally distinct macrophage subsets that can be identified in single-cell transcriptomics studies^20, 21^. To investigate more precisely how inhibiting GPIIb/IIIa-dependent thrombo-inflammation might modulate cardiac immune responses after MI, we performed single-cell RNA-sequencing (scRNA-seq) analysis of cardiac leukocytes with cell surface marker analysis by CITE-seq^24^ on day 4 after permanent MI in mice treated or not with JON/A-F(ab′)_2_ (**Figure 4a**). Biological replicates were included using cell hashing^25^. The total number of CD45^+^ cells did not differ between groups (**Figure 4b**). Clustering analysis (**Figure 4c-d**) and interrogation of cell surface markers (**Figure 4e**) and transcript expression (**Figure 4f**) allowed us to identify all immune cell lineages infiltrating the heart including neutrophil and macrophage subsets, overall corresponding to transcriptomic profiles described in previous studies^22,21^, including “young”, intermediate and SiglecF^+^ neutrophils, tissue-resident ‘TLF’ (Timd4^+^Lyve1^+^Folr2^+^) macrophages^26^, pro-inflammatory macrophages (*Il1b*), *Spp1* (encoding osteopontin) expressing macrophages, and macrophages with a ‘lipid associated macrophage’ (LAM) gene expression signature^21^. Subsets of type 1 and type 2 classical dendritic cells (cDC1, cDC2) and plasmacytoid dendritic cells (pDC), as well as T, B and NK cells were identified (**Figure 4c-d**). No differences in cellular distribution were observed (**Figure 4g-l**): proportion of total macrophages (**Figure 4g**) and distribution of macrophage subsets (**Figure 4h**), proportion of total neutrophils (**Figure 4i**) and distribution of neutrophil subsets (**Figure 4j**), proportion of dendritic cell subsets (**Figure 4k**), and proportions of lymphoid cells (**Figure 4l**) were not significantly altered between groups. Altogether, this analysis showed that inhibiting GPIIb/IIIa-dependent thrombo-inflammatory pathways-did not significantly impact acute cardiac inflammatory responses after MI.

**Figure 4:**
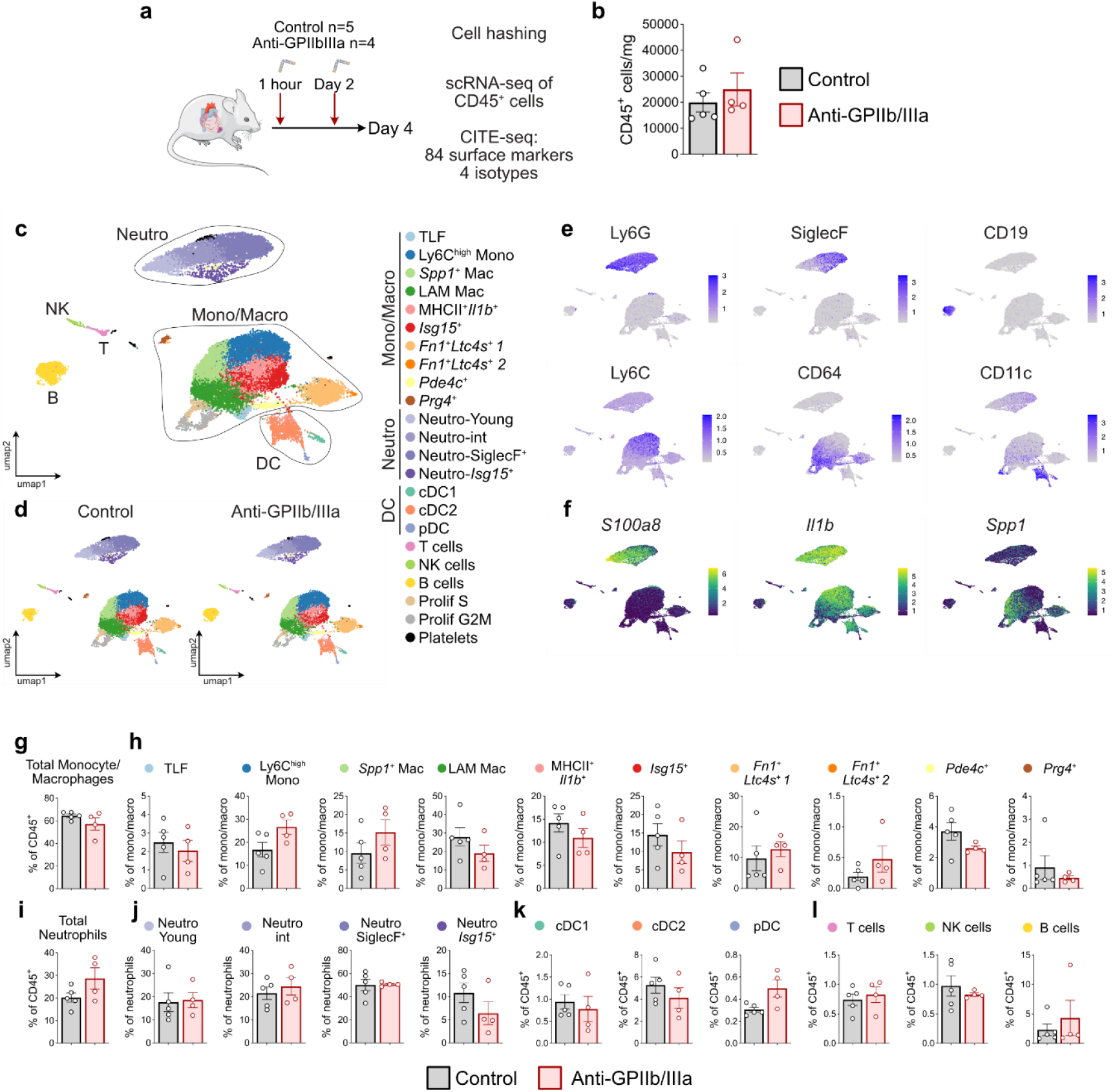
GPIIb/IIIa blockade and post-MI acute inflammatory responses: scRNA-seq. **a)** experimental workflow; **b)** Absolute counts of viable cardiac CD45^+^ cells. **c)** UMAP plot and identification of immune cell subsets; **d)** UMAP plot split according to experimental condition; **e)** expression of the indicated cell surface markers projected onto the UMAP plot; **f)** expression of the indicated transcripts projected onto the UMAP plot; **g)** proportion of total monocytes/macrophages among total cardiac CD45^+^ immune cells; **h)** proportion of monocyte/macrophage subsets among total macrophages; **i)** proportion of total neutrophils among total cardiac CD45^+^ immune cells; **j)** proportion of neutrophil subsets among total neutrophils; **k)** proportion of dendritic cell (DC) subsets among total cardiac CD45^+^ immune cells; **l)** proportion lymphoid cell subsets among total cardiac CD45^+^ immune cells. TLF: Timd4^+^Lyve1^+^Folr2^+^; LAM: lipid-associated macrophage; Neutro: neutrophil; Mono/Macro=monocyte/macrophage; cDC: classical dendritic cell; pDC: plasmacytoid dendritic cells; NK: natural killer. Panels g-l: data presented as mean ± SEM.

*Blockade of GPIIb/IIIa minimally affects post-MI cardiac repair*

We next analyzed survival and cardiac repair after permanent MI in JON/A-F(ab’)_2_ and control treated mice (**Figure 5a**). We observed a reduced survival in JON/A-F(ab’)_2_ -treated mice (**Figure 5b**). Paradoxically, no differences in infarct size (**Figure 5c-d**) or border zone fibrosis (**Figure 5e-f**) were observed. Altogether, these results indicate that while GPIIb/IIIa targeting in I/R injury might be beneficial, it does not ameliorate cardiac repair after permanent MI, but may instead be harmful, again underlining the impact of reperfusion as a major determinant of platelet-driven thrombo-inflammation.

**Figure 5:**
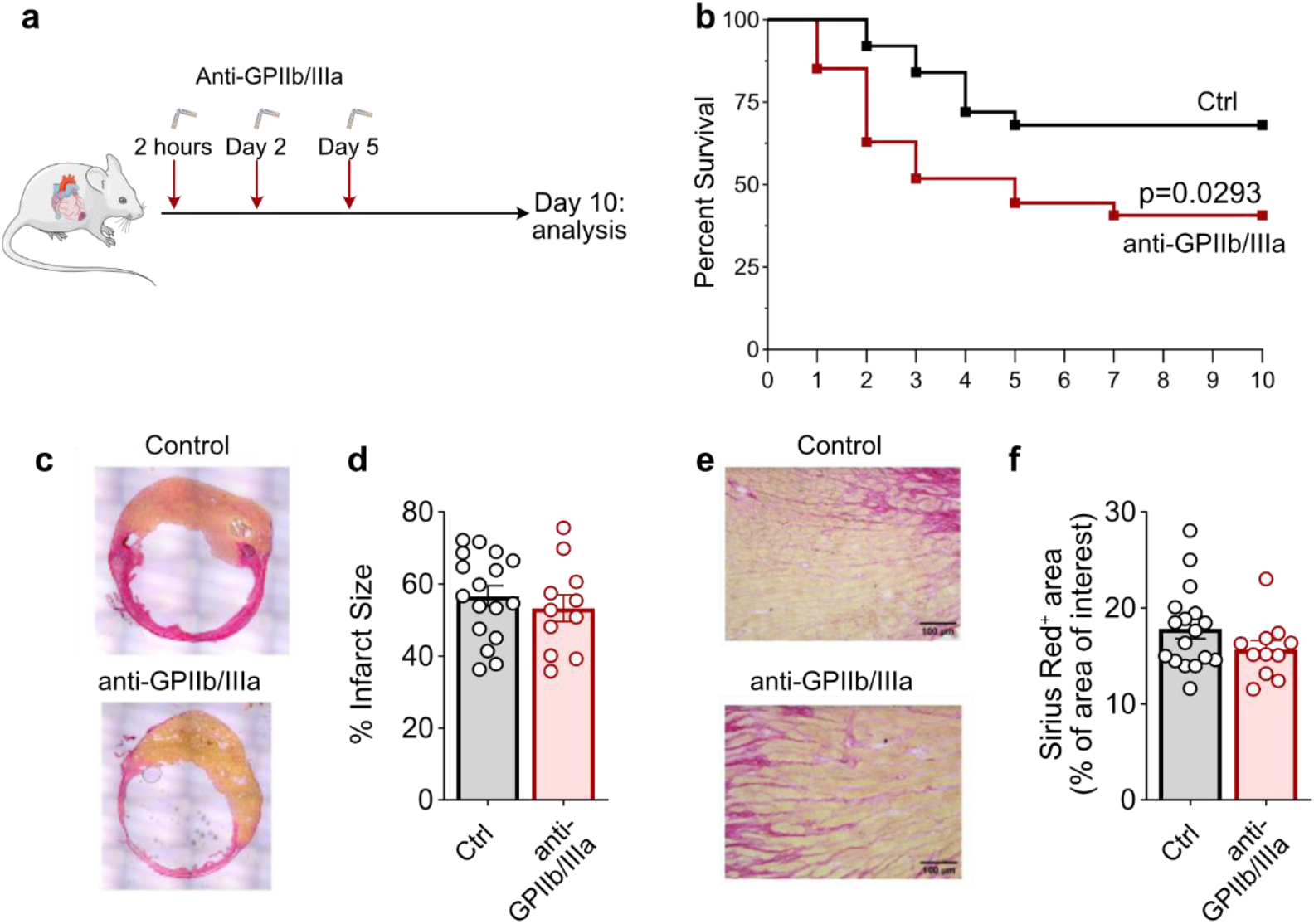
GPIIb/IIIa blockade and post-MI cardiac repair. **a)** experimental workflow; **b)** Survival plot of isotype control (n=25) and anti-GPIIb/IIIa treated (n=27) mice over 10 days after MI; **c)** Representative picrosirius red staining of heart sections and **d)** quantitative analysis of infarct size at day 10 after MI. **e)** Representative staining of picrosirius red heart sections and **f)** quantitative analysis of picrosirius red positive staining in the border zone. Scale bars: 100μm. Results pooled from 4 experimental series. **c-f)** control mice: n =17; JON/A treated mice: n =11. Data presented as mean ±SEM. Panel b: log-rank (Mantel-Cox) test; panel d and f: Mann-Whitney test.

## Discussion

The concerted interplay of platelets, leukocytes, endothelial cells and the coagulation cascade - a pathomechanistic principle referred to as thrombo-inflammation-is a powerful driver of myocardial damage after MI^8^. In previous studies, we have demonstrated that targeting thrombo-inflammation by blockade or depletion of GPVI is a promising approach to improve the (functional) outcome in ischemic conditions such as MI and stroke without further increasing bleeding risk^27^. In this study, we assessed the impact of GPVI blockade in a model of permanent ischemia. We depleted GPVI from the platelet surface using the JAQ1 IgG antibody and blocked GPIIb/IIIa using the JON/A F(ab’)_2_ fragment, and studied the effects on both inflammation (day 4) and cardiac repair (day 10) after MI.

In stark contrast to I/R models, GPVI or GPIIb/IIIa blockade did not substantially affect inflammatory responses, and only minimally impacted cardiac repair in permanent ischemia. This difference may be explained by the distinct pathophysiological mechanisms in the two models. In I/R, oxygenated blood can re-enter the ischemic myocardium enabling oxidative phosphorylation and pH normalization^28^. Along with erythrocytes, immune cells such as neutrophils and monocytes infiltrate the reperfused tissue, where they contribute to remodeling by clearing necrotic cells and initiating healing processes that culminate in the formation of a collagen-rich scar. However, in a Janus-faced process reperfusion can aggravate myocardial damage in reperfusion injury, which was demonstrated to contribute to up to 50% of the final infarct size and to play a central role in the pathophysiology of ischemic cardiomyopathy and lethal reperfusion arrhythmias^29^. Importantly, this “second wave” of inflammatory injury is entirely absent in a permanent ligation model. Due to the lack of collateral vasculature in the myocardium, the infarcted territory remains unperfused after permanent ligation, limiting immune cell influx^30^. Consistent with this concept, prior studies have demonstrated that permanent occlusion and I/R models exhibit distinct inflammatory kinetics, which are characterized by faster leukocyte accumulation and clearance in the reperfused myocardium^31^. Our findings suggest that the beneficial effect of anti-GPVI treatment is facilitated through the modulation of I/R processes. The mechanism by which GPVI impacts I/R remains incompletely understood.

The immunoreceptor GPVI is the central activating platelet receptor for collagen and fibrin(ogen)^32,33^. Besides its impact on platelet activation, aggregation and thrombus formation, blockade or depletion of GPVI was shown to reduce the infiltration of immune cell subsets in other models of inflammation such as acute respiratory distress syndrome, *Klebsiella pneumonia* induced infection, or zymosan-induced inflammation and mechanical allodynia^18, 34,35^. Nonetheless, our data show that in permanent ischemia GPVI depletion did not affect immune cell recruitment in the heart on day 4 after MI. Furthermore, loss of GPVI did not alter circulating leukocyte subset fractions and myelopoiesis in the bone marrow. Interestingly, however, GPVI-depleted mice displayed less interstitial fibrosis in the infarct border zone 10 days after MI. Thus, platelet GPVI might modulate collagen deposition, possibly mediated by platelet interaction with tissue repair promoting macrophages or with collagen-producing myofibroblasts, indicating a potentially increased left ventricular function after GPVI depletion post MI. Future studies focusing on cardiac function by using transthoracic echocardiography or MRI at later time points (e.g. 14 or 28 days after MI) are required to address this question.

Over the last decades, several therapeutics targeting GPVI have been evaluated comprising EMA601, glenzocimab and the competitive inhibitor revacept. In addition to the choice of the GPVI-targeting agent, also the clinical course of MI may be critical for applying GPVI blockade. Our study suggests that the benefit of GPVI inhibition is less pronounced in the setting of permanent occlusion, reflecting the substantial proportion of up to 28% of MI patients without sufficient recanalization in comparison to patients who achieve reperfusion^17,36^. These findings suggest that early application of GPVI inhibitors prior to reperfusion might be beneficial. To gain further translational insights, we have generated a mouse model expressing the human GPVI receptor (*hGP6*^*tg/tg*^), which enables us to study the impact of human GPVI inhibitors in vivo^37,11^. Further clinical studies are needed to define the optimal timing and patient subsets for GPVI-targeted therapy in MI.

We have shown in a previous study that platelets can actively drive macrophage activation and gene expression towards a pro-inflammatory phenotype in an in vitro co-incubation model^38^, which is reflected by upregulation of genes associated with inflammation such as *Il1b, Trem1, Tlr2, Cd14*. In our in vivo studies we did not detect differences in gene expression profiles after GPIIb/IIIa blockade. However, anti-GPIIb/IIIa treated mice displayed higher mortality compared to controls, suggesting that the worsened outcome after GPIIb/IIIa blockade was not caused by an altered leukocyte population. In addition, blockade of GPIIb/IIIa had no effect on infarct size and fibrosis. One explanation for the increased mortality could be increased bleeding, a known limitation for the use of GPIIb/IIIa inhibitors in the clinical setting, restricting their use mostly to bailout situations^27^. Our results are in accordance with the literature: blocking of platelet aggregation with the same anti-GPIIb/IIIa in a mouse model of myocardial I/R injury did not alter infarct size and immune cell recruitment^10^, while the same treatment in a tMCAO stroke model increased intracranial bleeding without affecting infarct size^39^.

In conclusion, our work demonstrates that the beneficial effect of targeting GPVI or GPIIb/IIIa might be of particular relevance for cardioprotection after I/R injury, whereas inflammation and tissue repair in permanent experimental MI may be operative mainly independently of platelet-driven pathways.

## Materials and Methods

### Mice

C57BL6/J male and female mice were purchased from Janvier lab (France). Mice were housed in 12-hour light/dark cycle between 7 am and 7 pm. All animal studies conform to the Directive 2010/63/EU of European Parliament and have been approved by the Government of Lower Franconia (Wuerzburg, Germany, Akt. 2-839, 2-743 and 2-865).

### Myocardial infarction and treatments

C57BL6/J male and female mice aged between 8 and 12 weeks were subjected to myocardial infarction by permanent ligation of the *left anterior descending* (LAD) coronary artery. 30 min before surgery, buprenorphin (0.1 mg/kg *subcutaneous* (s.c.)) was given to the mice for analgesia. Mice were anesthetized with isoflurane (induction 4%, maintenance 1.5-2.0%) and intubated using an endotracheal cannula, and placed on a heating pad under mechanical ventilation (Physitemp Instruments inc., TCAT-2LV or VentElite, Harvard apparatus). To confirm adequate anesthesia, the paw withdrawal reflex was checked. An ointment (Bepanthen 08109240) was applied on the eyes to protect them during the surgical procedure. The thorax area was shaved and disinfected. Skin was incised, the rib cage was revealed and a thoracotomy in the fourth intercostal space exposed the heart. With a 7/0 non-resorbable nylon suture thread (Serapren, CP05341A), permanent ligation of the left anterior descending artery was achieved. The thorax was closed by four independent sutures, while skin was closed with a continuous suture, using a 6/0 non-resorbable nylon suture thread (Serapren, CP07281A). Mice were administered buprenorphine as analgesic (0.1 mg/kg, s.c.) twice daily for the following two days after surgery, while Hydrogel (MediGel® Sucralose, ClearH2O) supplemented with buprenorphin (0.5 μg/mL buprenorphin in 60 mL MediGel® Sucralose, ClearH2O) was provided in the cage during the night for additional analgesia. Platelet GPIIb/IIIa was blocked by administering the mice with 4H5 i.e. Fab_2_ fragments of the JON/A antibody i.v 1 hour, 48 hours (day 2), and 120 hours (day 5) after LAD ligation at a dose of 100 μg/mouse. In both strategies, control mice were injected with the same volume of sterile PBS. To deplete platelet GPVI, mice were given the IgG JAQ1 antibody intraperitoneally (i.p) at a dose of 100 μg/mouse, on days 6, 4 and 2 before surgery as well as on day 4 after surgery, while control mice were treated with isotype control IgG. At the end of experiments, mice were euthanized by cervical dislocation under isoflurane anesthesia.

### Heart histology

After euthanasia, mice received an intracardiac perfusion of ice-cold PBS. The heart was excised, embedded in OCT compound (Tissue-Tek) and snap frozen in isopentane (Carl Roth, 3927.1) cooled in liquid nitrogen and stored at -80°C. 10 μm cryosections were collected with a Cryostat (Leica, CM3050 S), placed on SuperFrost slides (Langenbrick GmbH, 03-0060) and stored at -80°C until further analysis. Heart sections were stained following the picrosirius red solution manufacturer’s instructions. More specifically, slides were passed shortly in Xylene (VWR, 28975360) followed by decreasing concentrations of ethanol (Carl Roth, T913.3). Sections were washed twice in distilled water and then incubated for 1 hour in Picrosirius red (Sigma Aldrich, Direct Red 80, 365548-5G). Afterwards, they were washed briefly twice in acidified water (0.5% glacial Acetic Acid (Sigma-Aldrich, 33209) in water), then were dehydrated in increasing ethanol concentrations with a final wash in xylene. Mounting medium was Vectashield (Vectashield, H-5000). Images were acquired either with Thunder microscope (infarct size measurement) or Leica DM 4000 B LED microscope (fibrosis measurement). We performed image analysis with ImageJ software (Fiji).

### Flow Cytometry

After euthanasia, the heart was thoroughly perfused with ice-cold PBS, the left ventricle (infarct area and border zone) excised, minced and digested for 1 hour at 37°C using a thermomixer (Eppendorf, Thermomixer C) in an enzymatic digestion mix containing collagenase I (450 U/ml, Sigma-Aldrich, C0130), collagenase XI (125 U/ml, Sigma-Aldrich, C7657), hyaluronidase (60 U/ml, Sigma-Aldrich, H3506) and DNase I (60 U/ml, Sigma-Aldrich, DN25). The resulting cell suspension was filtered through a 70 µm cell strainer (Greiner bio-one, 542070), centrifuged at 400 rcf for 5 minutes at 4°C and resuspended in PBS containing 1% fetal calf serum (FCS).

Blood was collected retro-orbitally under deep anesthesia into EDTA coated tubes. 150 μL of blood was then mixed with 850 μL erythrocyte lysis buffer, centrifuged at 400 x g for 10 minutes at 4°C, and resuspended in PBS +1% FCS. One femur from each mouse was collected and bone marrow cells were extracted by centrifuging the bones for 1 minute at 10,000 rcf as described in^40^. The cell pellet was resuspended in in PBS +1% FCS.

Single cell suspensions of heart, blood and bone marrow were plated in a 96 round-bottom well plate (Sarstedt, 82.1582) followed by a centrifugation at 400 rcf at 4^°^C for 5 minutes. To block unspecific staining, samples were resuspended in TruStain FcX (Biolegend, 101320, working concentration= 1 μg/mL) and incubated for 15 minutes at 4^°^C in the dark. Next, cells were stained for 30 minutes at 4°C with the antibodies listed in table 1 and Fixable Viability Dye e780 (ThermoFisher 65-0865-14, 1:1000). 100 μL of counting beads (Biolegend, 424902) were added to the heart samples and the absolute cell number was calculated according to the manufacturer’s protocol. The samples were measured on a BD FACS Celesta (BD biosciences) and the analysis was performed with FlowJo v10 software. The pre-gating strategy for bone marrow, blood and heart cells is presented in **Methods Figure 1**.

**Table 1:** Flow Cytometry antibodies.

| Antibody | Clone | Catalog Number | Company | Final concentration (µg/mL) |
| --- | --- | --- | --- | --- |
| CD45 Alexa700 | 30-F11 | 103128 | Biolegend | 2.50 |
| CD45.2 BV421 | 104 | 109832 | Biolegend | 1.33 |
| CD45.2 APC | 104 | 109814 | Biolegend | 1.33 |
| CD11b BV510 | M1/70 | 101263 | Biolegend | 1.33 |
| CD11b PercpCy5.5 | M1/70 | 101228 | Biolegend | 1.33 |
| Ly6G Pacific Blue | 1A8 | 127612 | Biolegend | 2.50 |
| Ly6G Alexa700 | 1A8 | 127622 | Biolegend | 2.00 |
| Ly6C BV510 | HK1.4 | 128033 | Biolegend | 1.00 |
| Ly6C Alexa488 | HK1.4 | 128022 | Biolegend | 1.25 |
| CD115 APC | AFS98 | 135510 | Biolegend | 1.33 |
| CD115 PE | AFS98 | 12-1152-81 | ThermoFisher | 1.33 |
| F4/80 PECy7 | BM8 | 123114 | Biolegend | 8.00 |
| MHCII BV785 | M5/114.15.2 | 107645 | Biolegend | 1.33 |
| TIM4 Percpefluor710 | 54(RMT4-54) | 46-5866-80 | ThermoFisher | 4.00 |
| TIM4 PE | 54(RMT4-54) | 12-5866-82 | ThermoFisher | 4.00 |
| SiglecF PE | 1RNM44N | 12-1702-82 | ThermoFisher | 4.00 |
| SiglecF Alexa 700 | 1RNM44N | 56-1702-82 | ThermoFisher | 4.00 |

**Methods Figure 1 (related to Figure 1):**
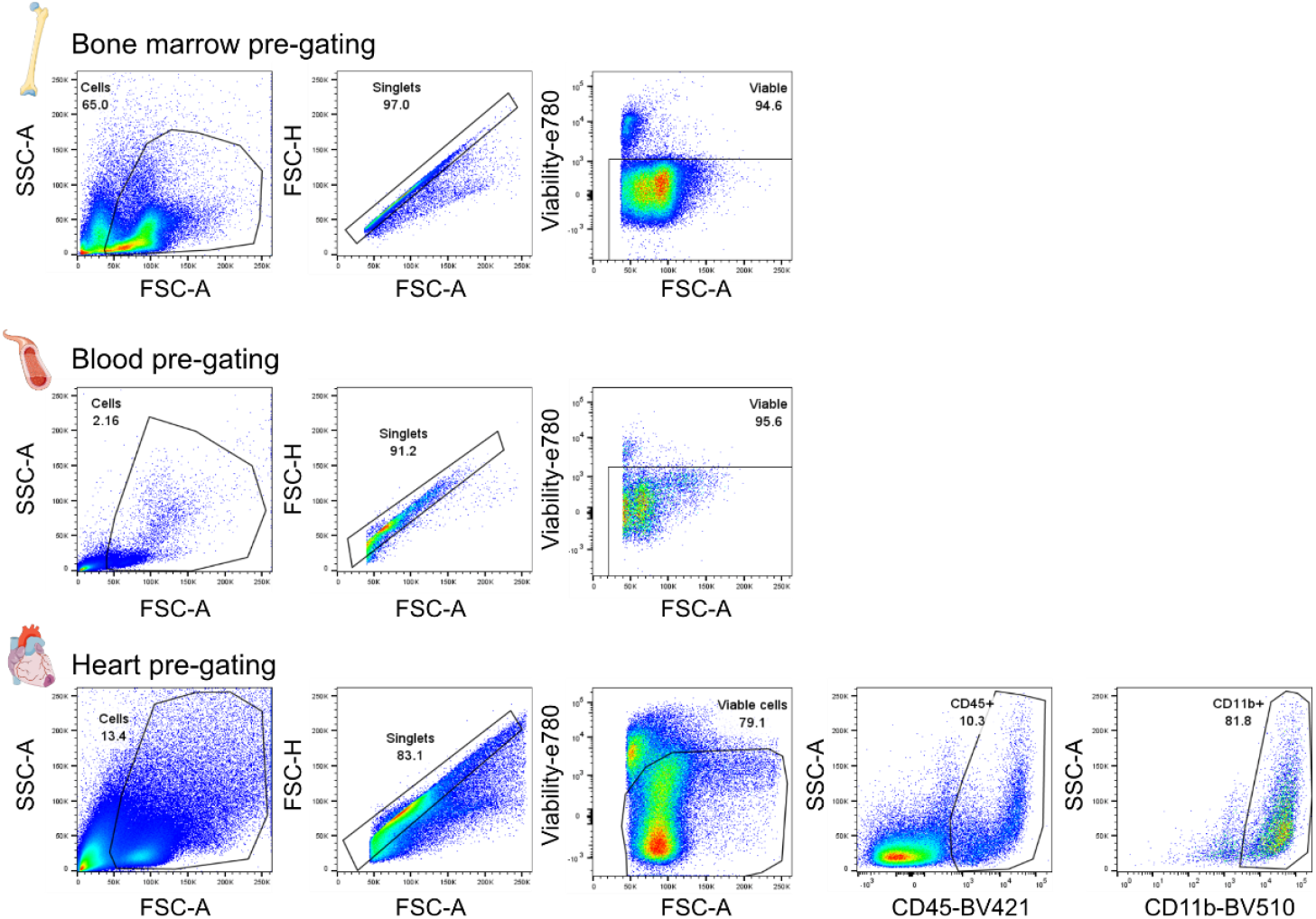
pre-gating strategy for bone marrow, blood and heart immune cells.

### Analysis of platelet glycoprotein receptor expression by flow cytometry

For analysis of platelet glycoprotein expression and activation, 50 μL of blood were collected 1.5 mL tubes containing Heparin. The diluted samples were stained with 10 μL of the following antibodies for 15 minutes at room temperature in the dark: active anti-GPIIb/IIIa-PE (in-house generated, clone JON/A), resting anti-GPIIb/IIIa-FITC (in-house generated, clone JON/A), anti-GPVI-FITC (in-house generated, clone JAQ1). The reaction was terminated by adding PBS and the samples were then measured by flow cytometry.

### Single-cell RNA-sequencing

The mice used in this experiment were all C57BL6/J on day 4 after MI, either treated with PBS (n=5) or with the JON/A F(ab’)_2_ anti-GPIIb/IIIa antibody 4H5 (n=4). Mice were anesthetized and injected i.v 5 minutes before sacrifice with 2 μg of anti-CD45.2 APC (Biolegend, clone 104, 109814) to label all circulating leukocytes. Mice were euthanized, hearts were flushed with ice-cold PBS, excised and the right ventricle was removed. Heart weight was measured and the hearts were minced with a scissor in Eppendorf tubes containing 500 μL of RPMI. These tubes were then incubated with an enzyme digestion mix containing collagenase I (450 U/mL, Sigma-Aldrich, C0130), collagenase XI (125 U/mL, Sigma-Aldrich, C7657), hyaluronidase (60 U/mL, Sigma-Aldrich, H3506) and DNase I (60 U/mL, Sigma-Aldrich, DN25) in a thermomixer (Eppendorf, Thermomixer C) for 45 minutes at 37°C with agitation. Afterwards, the cell suspension was filtered through a 70 μm cell strainer with undigested pieces being dissociated with a syringe plunger. Then, cells were washed in 25 mL of ice-cold PBS containing 1%FCS. After washing, samples were resuspended in 170 µL MACS Buffer (2 mM EDTA, 0.5% BSA in 500 mL PBS) containing 1:50 Fc Block and incubated for 5 to 10 minutes on ice. Afterwards, 30 µL MACS/FACS mix (24 µL CD45 microbeads (Militenyi, 130-052-301), 4 µL MACS-Buffer and 2 µL CD45.2 Alexa-488 (Biolegend, clone 104, 109815) were added in each sample for 15 minutes at 4^°^C. 1 µL of the corresponding hashtag antibody were added to each sample (BioLegend TotalSeq-A antibodies 1 to 9; Hashtag 1 to 5: PBS treated mice, hashtag 6 to 9: JON/A Fab_2_ anti-GPIIb/IIIa antibody) for another 15 minutes. Two washing steps followed, and cells from all samples were pooled. Positive magnetic selection of CD45^+^ cells was performed using LS columns and a QuadroMACS magnet (Militenyi Biotec) according to the manufacturer’s instructions. CD45^+^ cells were collected, washed twice and resuspended in PBS+1%FCS. The samples were stained with CITE-seq antibodies (BioLegend TotalSeq-A antibodies, see clones, references and concentrations in **Table 2**) together with live/dead e780 stain, then washed and processed to sorting with FACS Aria III (BD Biosciences). Live CD45^+^CD45iv^-^ cells were sorted with a 100μm nozzle.

**Table 2:** BioLegend TotalSeq-A antibodies.

| Antibody | ref. | [µg/ml] |
| --- | --- | --- |
| Hashtag 1 | 155801 | 2.5 |
| Hashtag 2 | 155803 | 2.5 |
| Hashtag 3 | 155805 | 2.5 |
| Hashtag 4 | 155809 | 2.5 |
| Hashtag 5 | 155811 | 2.5 |
| Hashtag 6 | 155813 | 2.5 |
| Hashtag 7 | 155815 | 2.5 |
| Hashtag 8 | 155817 | 2.5 |
| Hashtag 9 | 155819 | 2.5 |
| Hashtag 10 | 155821 | 2.5 |
| Hashtag 11 | 155823 | 2.5 |
| Hashtag 12 | 155825 | 2.5 |
| Ly6G | 127655 | 2 |
| CD11b | 101265 | 2.5 |
| CD62L | 104451 | 2 |
| IA/IE (MHCII) | 107653 | 1.67 |
| CD54 (ICAM1) | 116127 | 2.5 |
| Ly6C | 128047 | 1.25 |
| CD115 | 135533 | 2.5 |
| CXCR4 | 146520 | 2.5 |
| Msr1 | 154703 | 2.5 |
| CD64 | 139325 | 2.5 |
| FCeRIa | 134333 | 2.5 |
| CCR3 | 144523 | 2.5 |
| CD49d | 103623 | 2.5 |
| CD80 | 104745 | 2.5 |
| CD117 | 105843 | 2.5 |
| Sca1 | 108147 | 2.5 |
| CD11c | 117355 | 2.5 |
| TIM4 | 130011 | 2.5 |
| CX3CR1 | 149041 | 2.5 |
| XCR1 | 148227 | 2.5 |
| F4/80 | 123153 | 2.5 |
| CD86 | 105047 | 2.5 |
| CD135 | 135316 | 2.5 |
| CD103 | 121437 | 2.5 |
| CD169 | 142425 | 2.5 |
| CD8a | 100773 | 2.5 |
| SiglecH | 129615 | 2.5 |
| CD19 | 115559 | 2.5 |
| CD3 | 100251 | 2.5 |
| CD63 | 143915 | 2.5 |
| CD9 | 124819 | 2.5 |
| CD163 | 155303 | 2.5 |
| NK1.1 | 108755 | 2.5 |
| CD279 | 109123 | 2.5 |
| CD127 | 135045 | 2.5 |
| CD68 | 137031 | 2.5 |
| Sirpa | 144033 | 2.5 |
| CD274 | 153604 | 2.5 |
| ITGB7 | 321227 | 2.5 |
| CD26 | 137811 | 2.5 |
| MGL2 | 146817 | 2.5 |
| TCRgd | 118137 | 2.5 |
| CCR2 | 150625 | 2.5 |
| CD44 | 103045 | 2.5 |
| CD21/35 | 123427 | 2.5 |
| CD43 | 143211 | 2.5 |
| Hamster IgG | 400973 | 2.5 |
| Rat IgG1, κ | 400459 | 2.5 |
| Rat IgG2a, κ | 400571 | 2.5 |
| Rat IgG2b, κ | 400673 | 2.5 |
| CD47 | 127535 | 2.5 |
| SiglecF | 155513 | 2.5 |
| CD137 | 106111 | 2.5 |
| CD36 | 102621 | 2.5 |
| CCR5 | 107019 | 2.5 |
| CD278 | 117409 | 2.5 |
| PIR-A_B | 144105 | 2.5 |
| CD5 | 100637 | 2.5 |
| CD304 | 145215 | 2.5 |
| CD40 | 124633 | 2.5 |
| CD14 | 123333 | 2.5 |
| CD95 | 152614 | 2.5 |
| CD300c_d | 148005 | 2.5 |
| IL1RL1 | 145317 | 2.5 |
| TCRbeta | 109247 | 2.5 |
| Mac2 | 125421 | 2.5 |
| CD137L | 107109 | 2.5 |
| CD178 | 106613 | 2.5 |
| CD55 | 131809 | 2.5 |
| CD226 | 128823 | 2.5 |
| TIGIT | 142115 | 2.5 |
| CD39 | 143813 | 2.5 |
| JAML | 128507 | 2.5 |
| CXCR5 | 145535 | 2.5 |
| MGL1 | 145611 | 2.5 |
| CD24 | 101841 | 2.5 |
| CD88 | 135817 | 2.5 |
| CD81 | 104917 | 2.5 |
| CD83 | 121519 | 2.5 |
| Podoplanin | 127427 | 2.5 |
| IgM | 406535 | 2.5 |
| TIM3 | 119729 | 2.5 |
| BTLA | 139113 | 2.5 |
| CD223 | 125229 | 2.5 |
| CD25 | 102055 | 2.5 |
| CD152 | 106325 | 2.5 |
| KLRG1 | 138431 | 2.5 |
| CD41 | 133937 | 2.5 |
| CD4 | 100569 | 2.5 |

The collected cells were washed, resuspended in PBS enriched with 0.04% ultrapure BSA (ThermoFisher AM2618) and counted before loading in the 10x Genomics Chromium. A total of 33,000 cells were loaded in a volume of 22 μL in two duplicate 10x reactions. Chromium Single Cell 3’ Reagent Kits (v3.1 Chemistry) were used. Sequencing was performed with S2 100bp flowcell with Novaseq 6000 platform (Illumina) and the reads for CITE-Seq/Hashing sample were allocated as follows: 5% for the hashtags, 10% for the ADTs and 85% for the mRNAs.

10x Genomics data, HTO and ADT libraries were demultiplexed using Cell Ranger software (version 7.0.1). Alignment and counting steps were performed with the Mouse GRCm38 reference genome. The -feature-ref flag of Cell Ranger software was used to generate a gene expression matrix counts alongside the expression of cell surface proteins. The obtained gene-barcode matrix was further analyzed using Seurat v4^41^. Demultiplexing was performed in Seurat v4 to identify sample of origin, exclude multiplets, and cells with undetectable hashtag signal. Clustering analysis was performed using a standard Seurat workflow based on RNA levels. Briefly, low quality cells were removed (with more than 5% unique molecular identifiers (UMIs) mapping to mitochondrial transcripts), data were normalized, and the “vst” method was used to identify 2000 variable features. Afterwards, the “ScaleData” function was applied, principal component analysis (PCA) was performed. 20 PCs were used to perform clustering analysis and Uniform Manifold Approximation and Projection (UMAP) dimensional reduction and clustering at a 0.8 resolution. Immune cell lineages were identified using known surface markers of T cells (CD3), NK cells (NK1.1), B cells (CD19), dendritic cells (DCs: MHCII, CD11c) including XCR1+ cDC1, plasmacytoid DCs (SiglecH), neutrophils (Ly6G), F4/80^+^CD64^+^CX3CR1^+^Ly6C^low^ macrophages comprising both resident macrophages and macrophages appearing after MI, and monocytes (Ly6C).

### Statistical analysis

Statistical analysis was performed using GraphPad Prism 10.3.0. Normal distribution of data was first tested using a D’Agostino-Pearson test. Normally distributed data were analyzed using a parametric test (two-tailed t test), while non-normally distributed data were analyzed using a Mann-Whitney U test. Survival (Kaplan-Meier survival curves) was analyzed using a Log-rank (Mantel-Cox) test. All data are expressed as mean ± SEM. P values<0.05 were considered statistically significant.

## Acknowledgements

We thank the Single-Cell Center Würzburg for excellent technical support.

## Sources of Funding

This work was supported by the Deutsche Forschungsgemeinschaft (DFG) SFB1525 project number 453989101.

## Conflict of Interest

None.

## Data availability

Single-cell RNA-sequencing data generated for this paper will be made available upon publication.

## Notes

### Competing Interest Statement

The authors have declared no competing interest.

